# Toti: an integrated multi-omics database to decipher the epigenetic regulation of gene expression in totipotent stem cells

**DOI:** 10.1101/2025.01.24.634821

**Authors:** Yi Chai, Ruiying Zhang, Shunze Jia, Danfei Zhu, Siyi Chen, Xudong Fu, Xin Sheng

## Abstract

Totipotent cells (TSCs), the origin of mammalian life and foundation to early mammalian embryogenesis, are characterized by the highest differentiation capacity and extensive developmental potential. However, none of the existing embryonic databases have provided epigenetic and transcriptomic resources on totipotency, greatly limiting our understanding of the mechanisms governing the establishment and exit of totipotency. Here, we present Toti, a pioneering multi-omics database exclusively developed for totipotency, covering *in vivo*, *in vitro* and genome-edited human and mouse embryonic TSCs, TSC-like cells, pluripotent cells (PSCs), and embryos spanning preimplantation stages, with a total of 8,265 samples. Toti facilitates an in-depth exploration of the molecular mechanisms underlying totipotency by offering Search, Browse and Analysis modules available at http://toti.zju.edu.cn/.

## Introduction

Totipotent cells (TSCs), emerging in early embryogenesis with the broadest cellular plasticity in the mammalian body, harbor enormous potential for regenerative medicine and reproductive technology [1, 2]. Relative to the developmentally more restricted pluripotent cells (PSCs), TSCs are capable of producing all of the differentiated cell types in both embryo and extraembryonic components and forming an entire organism [3, 4]. Totipotency is limited to early-stage blastomeres. In mice, only the zygotes and blastomeres from 2-cell embryos are bona-fide TSCs and can give rise to the blastocyst, which is composed of the inner cell mass (ICM) and outer trophectoderm (TE) **(Figure 1A)** [5–7]. As cells develop into distinct cell lineages by the blastocyst stage, totipotency is lost and cellular plasticity is gradually reduced. Currently, due to the scarcity of embryonic TSCs, the gene regulatory logic underlying totipotency is still incompletely understood.

**Figure 1.**
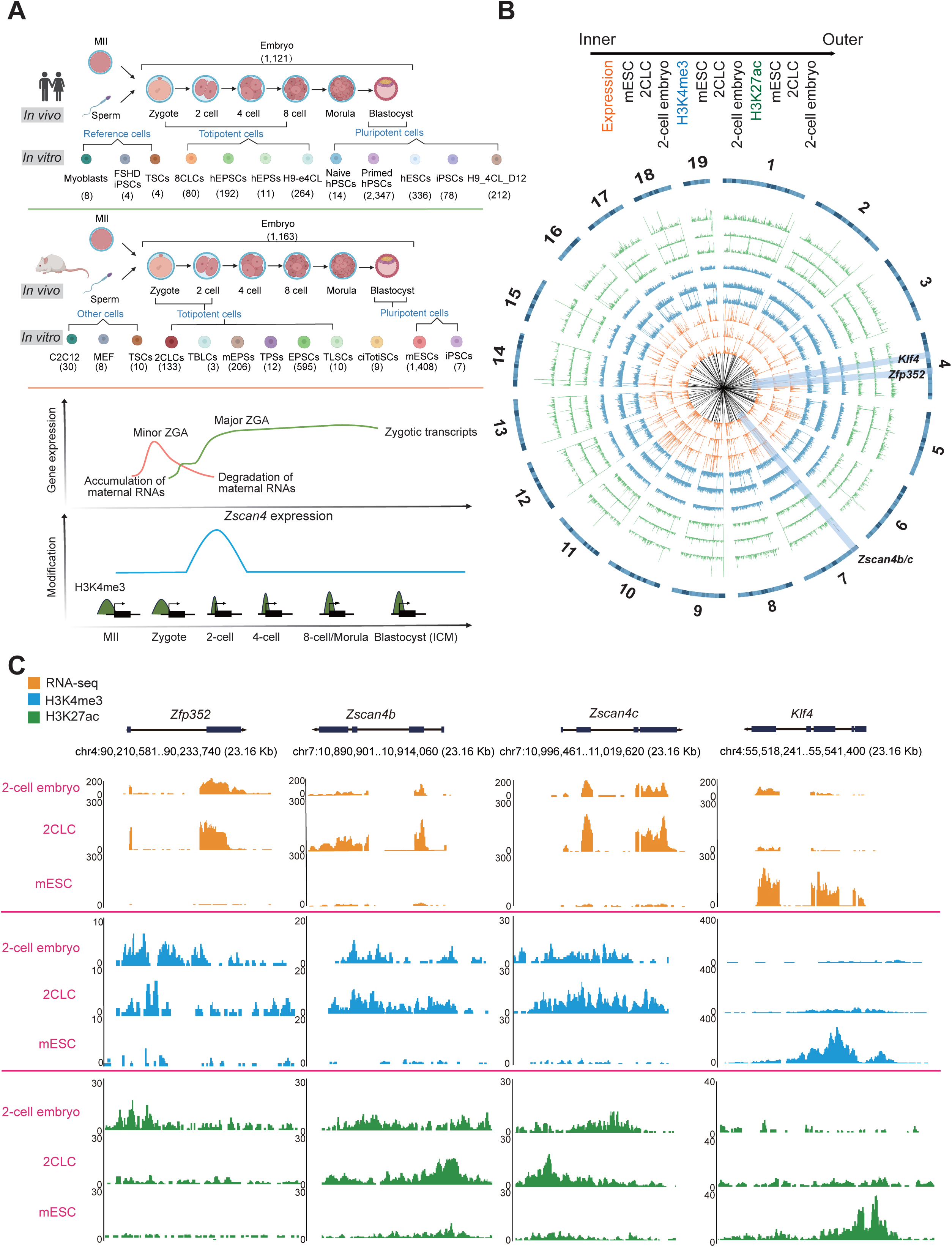
Toti provides multi-omics data to investigate epigenetic and transcriptomic underpinning of totipotency. **A.** Toti contains multi-omics data in *in vivo*, *in vitro* and genome-edited embryonic TSCs, TSC-like cells, PSCs and embryos spanning preimplantation stages from 4,671 human and 3,594 mouse samples. FSHD: Facioscapulohumeral Muscular Dystrophy, PSCs: pluripotent stem cells, TSCs: trophoblast stem cells, 8CLCs: 8-cell like cells, EPSCs: expanded pluripotent stem cells, EPSs: extended pluripotent stem cells, H9-e4CL: (H9 hESC cultured in e4CL (stepwise) for 5 days), ESCs: embryonic stem cells, C2C12: cultured mouse myoblasts, MEF: mouse embryonic fibroblasts, 2CLCs: 2-cell-like cells, TBLCs: totipotent blastomere-like cells, TPSs: totipotent potential stem cells, EPSCs: expanded pluripotent stem cells, TLSCs: totipotent-like stem cells, ciTotiSCs: chemically induced totipotent stem cells, ZGA: zygotic genome activation. **B.** A Ciros plot showing reads coverage for gene expression (orange rings), H3K4me3 (blue rings), and H3K27ac (green rings) histone modified peaks in mESC, 2CLC, and 2-cell embryo. The black lines in the center represent genomic gaps or low mappability regions, such as centromeres and other repetitive sequences. **C.** Reads coverage represents gene expression (orange), H3K4me3 (blue), and H3K27ac (green) modifications around totipotent signature genes, *Zfp352*, *Zscan4b*, and *Zscan4c*, and pluripotent signature gene *Klf4*, in 2-cell embryo, 2CLC, and mESC. 2CLC: 2-cell-like cell, mESC: mouse embryonic stem cell.

*In vitro* cellular models are critical for understanding the molecular architectures of cell stemness. For instance, embryonic stem cells (ESCs), derived from the ICM of blastocysts [3], are classic pluripotent cellular models and greatly facilitate the exploration of pluripotency [8]. Intriguingly, approximately 1-5% of ESCs resembling the blastomeres of 2-cell embryos, referred to as ‘2-cell-like-cells’ (2CLCs), arise spontaneously in ESC culture *in vitro* [9–11]. They show greater cellular plasticity and downregulated protein levels of pluripotency factors [9–11]. Unlike pluripotent ESCs, 2CLCs retain the totipotent-like state to generate both embryonic and extraembryonic tissues when reintroduced into early embryos [9], highlighting the promising application of these cells in understanding totipotency. Other *in vitro* totipotent-like cellular counterparts with biological relevance, including totipotent blastomere-like cells (TBLCs) [12], totipotent potential stem cells (TPSs) [13], and chemically induced totipotent stem cells (ciTotiSCs), have been actively pursued afterwards [14, 15] and widely used in totipotency study as well.

Recent advances in totipotency cellular model and sequencing technology lead to the ever-growing influx of multi-omics data on totipotent-like cells. These results have rapidly forwarded our understanding of epigenomic and transcriptomic features governing the establishment and exit of totipotency [16–24]. For example, previous studies have identified transcription factor *Dux* as the master inducer of the 2C-like transition in mESCs [16–18, 25, 26], based on RNA-seq and ChIP-seq data detected from CRISPR-based genome-edited embryonic cells. Supported by RNA-seq data, 2CLCs are typically characterized by transient activation of major satellites [19] and endogenous retroviral (ERV) elements that significantly contribute to promoting ZGA [4]. Particularly, as shown by comprehensive epigenetic information measured by ATAC-seq, Bisulfite sequencing, and ChIP-seq, endogenous retrovirus, MERVL, activated many 2C-specific genes, including the *Zscan4* cluster genes, likely through epigenetic variations, such as DNA methylation and histone modifications associated with DNMT [20, 25] and LSD1 (KDM1A) [21–23]. Other features of 2-cell embryos, such as their chromatin accessibility landscape [17] and increased global histone mobility [24], are also found to be recapitulated in 2CLCs. Moreover, the remodeling of the totipotency-specific broad H3K4me3 domains could help stabilize totipotency *in vitro* [27], highlighting the importance of characterizing epigenetic underpinning related to totipotency. Besides, benefiting from scRNA-seq data, our previous studies identified a novel intermediate state during the 2C-like transition [25, 26]. Therefore, extensive investigation of epigenetic and transcriptomic architectures in totipotent cells is of paramount importance to pinpoint molecular mechanisms controlling cellular plasticity and lineage segregation in early development.

Integrative analysis of multi-omics data in both embryonic cells and their *in vitro* counterparts makes it possible to comprehend the molecular basis of *in vivo* cell fate differentiation and *in vitro* cell fate engineering. Currently, arising databases mainly contain information on pluripotent cells instead of totipotent cells and few information on CRISPR-based genome-edited embryonic cells. For example, EMAGE [28], DBTMEE [29], EmExplorer [30], StemCellDB [31], LifeMap Discovery [32], FunGenES [33] and StemMapper [34] provide gene expression patterns during mouse or human embryo development and in induced pluripotent stem cells (iPSCs). ScRNASeqDB [35] provides single-cell gene expression profiles in 200 human early embryonic cells at different developmental stages. MethBank 3.0 [36], DevMouse [37], iHMS [38], GED [39], and MetaImprint [40] are mainly focused on epigenetic modifications, such as DNA methylation and histone modifications during embryogenesis in human and mouse. EpiDenovo [41], ESCAPE [42], DevOmics [43], and dbEmbryo [44] integrated genomic, epigenomic, and transcriptomic data from human and mouse early embryos without TSCs. However, none of the existing databases have provided multi-omics data on *in vivo*, *in vitro,* and genome-edited totipotent cells, greatly hindering the interpretation of underlying mechanisms that contribute to totipotency and embryo development.

To bridge this gap, here we present a novel database, Toti, the unique and first multi-omics database dedicated to comprehensively investigating epigenetic and transcriptomic architectures in early-stage embryogenesis of mouse and human **(Figure 1A).** Toti encompasses *in vivo*, *in vitro*, and genome-edited embryonic cells, with the particular focus on totipotent cells, thus providing an unprecedented platform for extensive investigation of totipotency *in silico* **(Figure 1A)**. Toti allows facilitated and quick access of interested samples/datasets by searching for genes, sequencing types or other relevant keywords. Toti enables comparative analyses not only on epigenetic characteristics measured by multiple methods, including bisulfite sequencing, ATAC-seq, ChIP-seq, CUT&RUN, and CUT&TAG, but also on temporal gene expression patterns supported by RNA-seq and scRNA-seq data to dissect molecular (dis)similarity across *in vivo* and *in vitro* totipotent cells, PSCs, and other embryonic cells in human and mouse **(Figure 1B)**. Toti also facilitates users to prioritize top enriched transcription factors (TFs), genes and pathways under different developmental stages and genome-edited conditions. Taken together, Toti thus serves as a unique, comprehensive, and valuable resource to provide insights into molecular characteristics shaping totipotency.

## Results

### Toti overview

As characterization of the molecular basis in TSCs is the key to comprehend mechanisms underlying totipotency, here we design Toti to store gene expression and epigenetic modification information covering *in vivo*, *in vitro* or CRISPR-based genome-edited totipotent, pluripotent, and differentiated embryonic cells **(Figure 1A)** from 4,671 human and 3,594 mouse samples **(Tables S1 and S2)**. Toti contains: (1) DNA methylation (WGBS, PBAT, and RRBS), (2) genome-wide chromatin features, including open chromatin peaks (ATAC-seq), TF binding sites, and histone modifications (ChIP-seq, CUT&TAG, and CUT&RUN), (3) gene expression patterns (RNA-seq, scRNA-seq) from different types of cells. According to these holistic multi-omics data in Toti, we observed globally similar patterns of peaks in gene expression, H3K4me3 and H3K27ac modifications in both mouse 2-cell embryos and 2CLCs, but distinct from mESCs (**Figure 1B**). Specifically, regarding totipotent genes, such as *Zfp352* [45], *Zscan4b,* and *Zscan4c* [46], enriched peaks in gene expression, H3K4me3, and H3K27ac were shown in both mouse 2-cell embryos and 2CLCs, but not in mESCs, highlighting their totipotency-specific signatures in 2-cell embryos and 2CLCs (**Figure 1C**). As a comparison, we also observed mESC-specific peaks in gene expression, H3K4me3, and H3K27ac around the pluripotent gene *Klf4* (**Figure 1C**) [47]. Hence, by integrative analysis of these orthogonal omics data from cells under different developmental stages or genome-edited conditions, Toti makes it possible for users to comprehensively investigate totipotency *in silico.* Toti is designed with three main functionalities: Search, Browse, and Analysis **(Figure 2)**.

**Figure 2.**
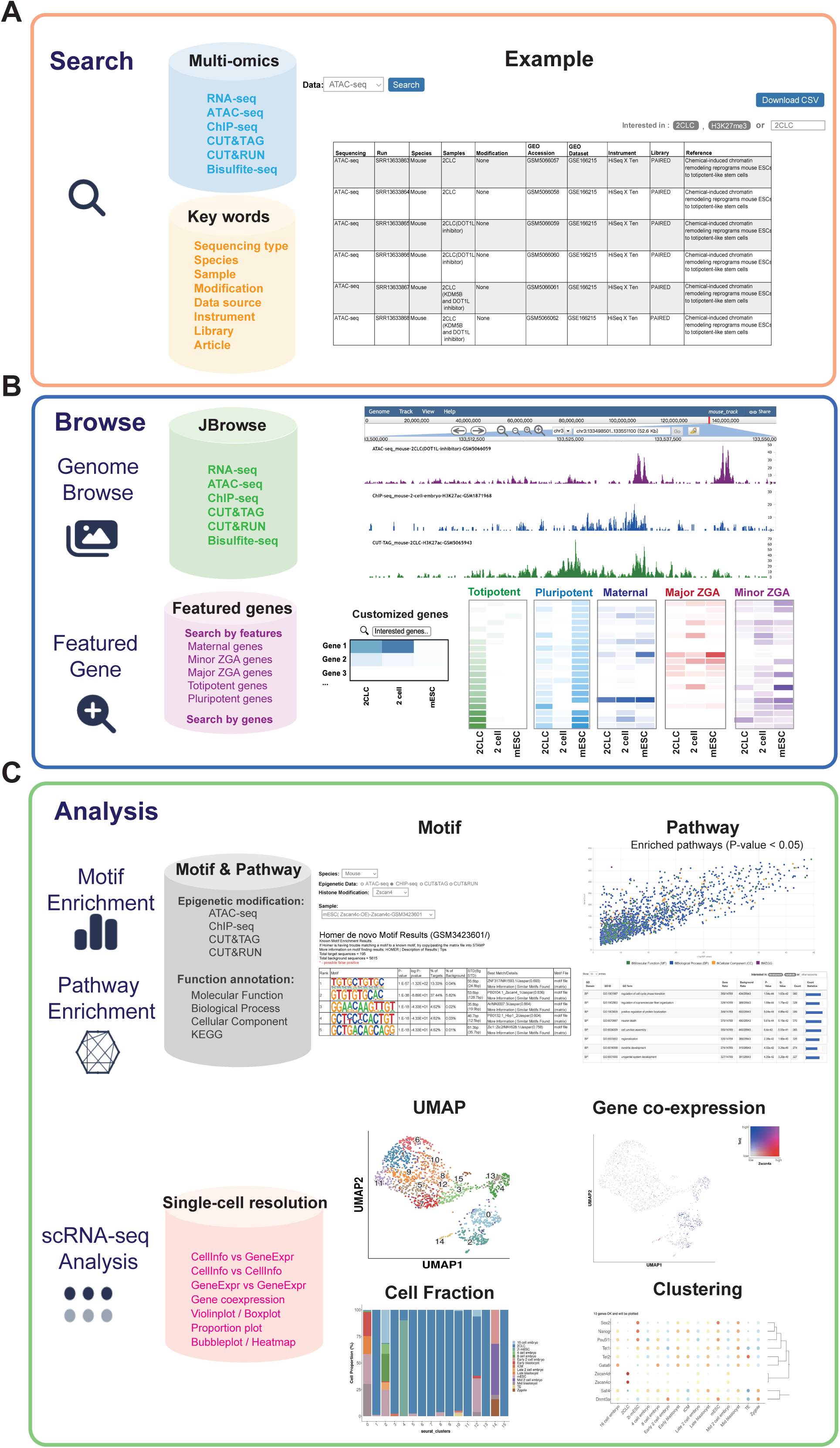
The main function of Toti. **A.** ‘Search’ module allows a flexible search of genes, sequencing types, or other relevant keywords. **B.** ‘Browse’ includes two sub-modules. ‘Genome Browse’ visualizes analyzed multi-omics data by embedding configured JBrowse. ‘Featured Gene’ visualizes expression patterns of top variable signature genes or user-selected genes under selected developmental stages or genome-edited conditions. ZGA: zygotic genome activation. **C.** ‘Analysis’ contains three sub-modules. ‘Motif Enrichment’ and ‘Pathway Enrichment’ prioritize enriched transcription factors and biological pathways associated with interested development stage or genome-edited condition, respectively. ‘scRNA-seq Analysis’ facilitates the intuitive comparison of gene expression or co-expression patterns among embryos under selected developmental stages or genome-edited conditions.

### Search

Toti enables a flexible search that provides quick access to totipotency-related studies/datasets of interest by querying genes, sequencing types, species, or other relevant keywords, such as histone modification type, cell type, library strategy, article name, and GEO accession ID. All the queried tables recording meta information of data stored in Toti are sortable and downloadable. Users can also click the hyperlink of each GEO accession ID to obtain further experimental details or acquire the corresponding raw data (**Figure 2A**).

### Browse

A systematic scrutiny of epigenetic and transcriptomic architectures, specifically for signature genes representing for totipotency or pluripotency, is extremely important to understand key factors governing the establishment and exit of totipotency in embryos. Toti ensures both orthogonally and parallelly comparative investigations on molecular basis with ‘Genome Browse’ and ‘Featured Genes’ **(Figure 2B)**.

### Genome Browse: allows for orthogonally and parallelly comparative analysis

Toti provides extensive investigation into gene expression and epigenetic variations, such as open chromatin peaks, histone modifications, and DNA methylation, across *in vivo*, *in vitro* and genome-edited totipotent, pluripotent, and differentiated cells by embedding our well-configured JBrowse **(Figure 2B)**. Data are hierarchically organized based on species, sequencing type, epigenetic modification, and gene expression. The interface permits the selection of one or multiple tracks to visualize analyzed multi-omics data. By specifying interested species, cell type, gene, or chromosome region, tracks for the corresponding regions are displayed. And it is of particular convenience to share the queried result to others for further discussion by clicking the ‘Share’ button on the top right of the Genome Browser. Users can also add custom data tracks for personalized comparison. All the resulting images, such as plots for gene structure, reads coverage, and histone modification peaks, are easily to be exported.

### Featured genes: provides insights into expression patterns of signature genes

The developmental trajectory of totipotent cells during early development is governed by highly coordinated transcriptomic remodeling. After fertilization, the maternal genes inherited from oocytes drive two-waves of zygotic genome activation (ZGA), including minor ZGA and major ZGA [48]. These ZGA genes modulate the entry and exit of totipotency in embryos [48]. Understanding the intricate interplay of these signature genes is critical for comprehending the biological nuances of totipotency. Here, we designed ‘Featured Genes’ **(Figure 2B)** sub-module to allow users to scrutinize expression patterns of signature genes, including maternal genes, minor ZGA genes, major ZGA genes, totipotent genes, and pluripotent genes defined from Yang et al. [27], across both *in vivo* and *in vitro* TSCs, and cells under different developmental stages or genome-edited circumstances. Users can customize their search by choosing interested featured genes and specifying cells to compare. Toti prioritizes development-or genome-edited-condition-associated genes based on the standard deviation of gene expression among selected samples. Subsequently, interactive heatmaps are rendered, providing a visual representation of top variable signature genes across chosen samples. Alternatively, user can also flexibly select any gene of interest and visualize their expression patterns across selected samples. The expression matrix of each heatmap can be downloaded and further tailored for customized use.

### Analysis

It is of fundamental importance to pinpoint top enriched transcription factors (TFs), genes, and pathways under different totipotent-associated development stages or genome-edited conditions, as these analyses could shed light on gene regulatory logics contributing to totipotency. Here, we designed three sub-modules, including ‘Motif Enrichment’, ‘Pathway Enrichment’, and ‘scRNA-seq analysis’ in this ‘Analysis’ section **(Figure 2C)** to provide insights into the top development-or genome-edited-condition-associated TFs, genes, or pathways, and facilitate the dissection of gene expression patterns at single-cell resolution in totipotency-related samples.

### Motif Enrichment: prioritizes top relevant transcription factors

As transcription factors play important roles in modulating cellular gene expression, we developed a ‘Motif Enrichment’ sub-module to prioritize top associated transcription factors (TFs) under different genome-edited conditions, histone modifications, or developmental stages **(Figure 2C)**. By selecting interested epigenetic modifications, such as open chromatin peaks (ATAC-seq), TF binding sites, or histone modifications (ChIP-seq, CUT&TAG, or CUT&RUN), in cells under certain conditions, Toti provides enriched TF-binding motifs analyzed by HOMER (http://homer.ucsd.edu/homer/). Further details for motifs are accessible via clicking links, such as ‘motif file’ and ‘Known Motif Enrichment Results’.

### Pathway Enrichment: prioritizes top associated biological pathways

Previous studies indicated that recapitulating epigenetic modifications of 2-cell embryos, such as chromatin accessibility landscape [17] and broad H3K4me3 [27], can facilitate totipotency stabilization *in vitro*. This underscores the importance of characterizing activated biological pathways under the establishment, self-renewal, or exit of totipotency. Therefore, we developed the ‘Pathway Enrichment’ sub-module **(Figure 2C)**, providing users with an intuitive and interactive chart, along with a sortable and downloadable table that prioritizes key relevant biological pathways based on gene ontology (GO) [49] and KEGG [50]. This sub-module enables users to gain insights into biological context dependent epigenetic modifications in *in vivo*, *in vitro* or genome-edited totipotent, pluripotent, and differentiated cells. Each resulting table can be sorted and downloaded. More information, such as genes involved in each pathway, could be accessible in the downloaded CSV file.

### scRNA-seq analysis: dissects transcriptome nuances at single-cell resolution

Using scRNA-seq data sequenced at different time points of *Dux* induction, we previously confirmed the existence of an intermediate state during the 2C-like transition [25], emphasizing that scrutiny of transcriptome at single-cell resolution could promote a better understanding of molecular mechanisms underlying the establishment and exit of totipotency. Here, we presented an interactive ‘scRNA-seq analysis’ interface to facilitate the intuitive comparison of gene expression or co-expression across cells under different developmental stages and TSC-like cells with/without genome-editing **(Figure 2C)**. To our knowledge, this is the first online scRNA-seq analysis tool tailored to investigation in transcriptome nuances of totipotency.

### A Case Study: a systematic integrated analysis of epigenetic and transcriptomic architectures in totipotency

To guide users through the multifaceted functionality and potential extension for Toti, here we present an integrative case study that delves deep into epigenetic and transcriptomic nuances of totipotency. In this case study, we mainly chose five mouse cell types for comparison, including pluripotent mESCs, embryonic TSCs (2-cell embryos), and three *in vitro* TSCs models (2CLCs, TPSs, and ciTotiSCs). The 2CLC is a natural transient totipotent-like cell population within mESC cultures [9–11], while TPS and ciTotiSC are two types of chemical-induced totipotent-like cell lines. All these three *in vitro* TSC models were previously found to show similar upregulated expression of totipotency factors as in totipotent 2-cell (2C) embryos but were distinguished from mESCs [10, 13, 51]. Our case study aims to delineate epigenetic and transcriptomic architectures recapitulated by these *in vitro* TSCs, respectively, through systematic integrated analysis of multi-omics data available in Toti. This case study not only reveals discrepancies in totipotency-associated molecular features within *in vitro* TSC models, but also sheds light on how to leverage these models in totipotent exploration. The modules in Toti we applied in this study include ‘Featured Gene’, ‘Genome Browse’, ‘Motif Enrichment’, ‘Pathway Enrichment’, ‘ScRNA-seq analysis’, and ‘Download’.

### Deciphering transcriptome discrepancies in totipotency across *in vitro* TSC models

The *in vitro* TSCs exhibit totipotent-like features in terms of transcriptome and developmental potential, and thus serve as alternative models for totipotency study [52]. To evaluate totipotency-associated characteristics these *in vitro* TSCs recapitulated, we chose embryos across different developmental stages to compare the molecular (dis)similarity across these cellular models and *in vivo* cells using scRNA-seq analysis. mESCs were also included in the analysis as controls. We observed that all *in vitro* TSCs showed higher similarity with the totipotent 2-cell embryos compared to mESCs **(Figure 3A)**. Interestingly, although these TSCs models were all considered totipotent-like cells, there were discrepancies within their gene expression patterns, as evidenced by their separated clustering positions in the UMAP **(Figure 3A)**.

**Figure 3.**
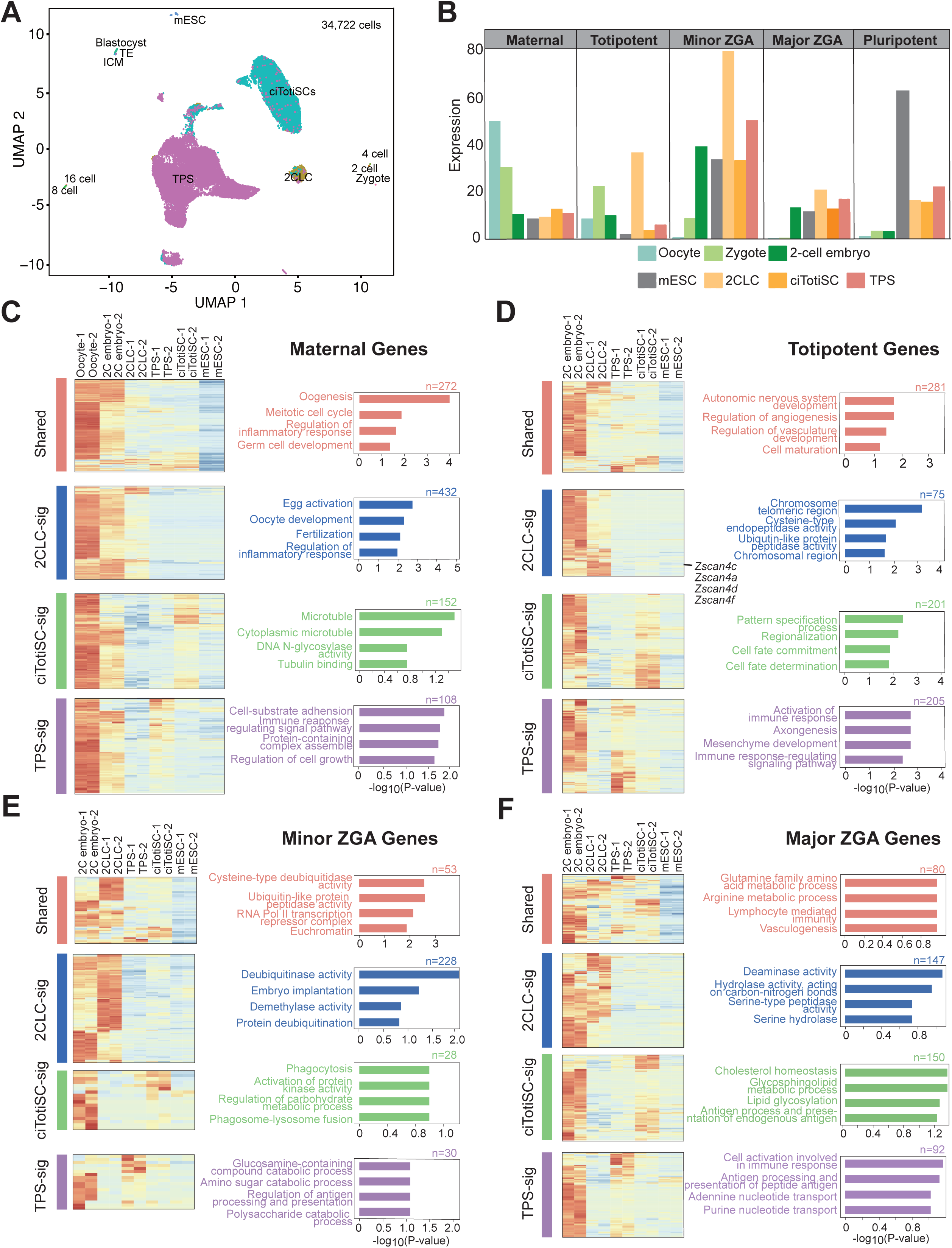
Transcriptome discrepancies in totipotency across *in vitro* TSC models. **A.** The UMAP of *in vitro* mouse TSC models (2CLCs, ciTotiSCs, and TPSs), mESCs and mouse embryo cells spanning from zygote to blastocyst (n = 34,722 cells). 12 cell clusters were identified. 2CLC: 2-cell-like cell, ciTotiSC: chemically induced totipotent stem cell, TPS: totipotent potential stem cell. **B.** Expression patterns of maternal genes, minor ZGA genes, major ZGA genes, totipotent genes, and pluripotent genes in oocyte, zygote, 2-cell embryo, mESC, 2CLC, ciTotiSC, and TPS. ZGA: zygotic genome activation, mESC: mouse embryonic stem cell, 2CLC: 2-cell-like cell, ciTotiSC: chemically induced totipotent stem cell, TPS: totipotent potential stem cell. **C.** Heatmaps showing gene expression patterns of maternal genes in oocytes, 2-cell embryos, 2CLCs, TPSs, ciTotiSCs and mESCs (left). Functional enrichment (gene ontology, GO) analysis results (right). The x axis is the -log_10_(two-sided P-value); The y axis represents the enriched pathways. Shared: genes expressed in all three *in vitro* TSC models (2CLCs, ciTotiSCs, and TPSs), 2CLC-sig/ciTotiSC-sig/TPS-sig: cell-type-specific expressed genes, 2CLCs: 2-cell-like cells, TPSs: totipotent potential stem cells, ciTotiSCs: chemically induced totipotent stem cells, mESCs: mouse embryonic stem cells. **D.** Heatmaps showing gene expression patterns of totipotent genes in 2-cell embryos, 2CLCs, TPSs, ciTotiSCs and mESCs (left). Functional enrichment (gene ontology, GO) analysis results (right). The x axis is the -log_10_(two-sided P-value); The y axis represents the enriched pathways. Shared: genes expressed in all three *in vitro* TSC models (2CLCs, ciTotiSCs, and TPSs), 2CLC-sig/ciTotiSC-sig/TPS-sig: cell-type-specific expressed genes, 2CLCs: 2-cell-like cells, TPSs: totipotent potential stem cells, ciTotiSCs: chemically induced totipotent stem cells, mESCs: mouse embryonic stem cells. **E.** Heatmaps showing gene expression patterns of minor ZGA genes in 2-cell embryos, 2CLCs, TPSs, ciTotiSCs and mESCs (left). Functional enrichment (gene ontology, GO) analysis results (right). The x axis is the -log_10_(two-sided P-value); The y axis represents the enriched pathways. Shared: genes expressed in all three *in vitro* TSC models (2CLCs, ciTotiSCs, and TPSs), 2CLC-sig/ciTotiSC-sig/TPS-sig: cell-type-specific expressed genes, 2CLCs: 2-cell-like cells, TPSs: totipotent potential stem cells, ciTotiSCs: chemically induced totipotent stem cells, mESCs: mouse embryonic stem cells. **F.** Heatmaps showing gene expression patterns of major ZGA genes in 2-cell embryos, 2CLCs, TPSs, ciTotiSCs and mESCs (left). Functional enrichment (gene ontology, GO) analysis results (right). The x axis is the -log_10_(two-sided P-value); The y axis represents the enriched pathways. Shared: genes expressed in all three *in vitro* TSC models (2CLCs, ciTotiSCs, and TPSs), 2CLC-sig/ciTotiSC-sig/TPS-sig: cell-type-specific expressed genes, 2CLCs: 2-cell-like cells, TPSs: totipotent potential stem cells, ciTotiSCs: chemically induced totipotent stem cells, mESCs: mouse embryonic stem cells.

To explore discrepancies across these TSC models, we used the ‘Featured Gene’ module to scrutinize expression patterns of maternal genes, minor ZGA genes, major ZGA genes, totipotent genes, and pluripotent genes in these TSC models. In line with previous studies, these TSC models exhibited elevated expression of totipotent and ZGA genes, and decreased expression of pluripotent genes compared to mESCs **(Figure 3B)** [52]. Notably, we also observed differences in expression of maternal and totipotent genes in these *in vitro* TSCs, while comparing to that in zygotes **(Figure 3B)**, suggesting the transcriptome diversity of captured totipotency in these *in vitro* models.

Next, we categorized maternal, totipotent, and ZGA genes into TSCs-shared cluster and cell-type-specific clusters, including 2CLC-signature cluster, cTotiSC-signature cluster, and TSC-signature cluster **(Figures 3C-3F, Table S3)** based on unsupervised hierarchical clustering on averaged expression levels across samples for each cell type using Euclidean distance and complete linkage method. Our analysis show distinct totipotent-associated characteristics for each *in vitro* TSC model **(Figures 3C-3F)**. For instance, 2CLCs exhibited increased expression in totipotent genes enriched in pathways associated with chromosome telomeric region compared to that of ciTotiSCs and TSCs **(Figure 3D)**, indicating 2CLCs may exhibit chromatin alterations relevant to totipotency. Notably, the well-known totipotent marker genes of *Zscan4* family are presented in this 2CLC-specific cluster, which is in line with previous investigations on both transcriptomic and epigenetic data in 2CLCs [20, 53]. Moreover, we also observed that cTotiSC-signature totipotent genes showed enrichment in cell-fate commitment, pattern specification, and regionalization **(Figure 3D)**. TPSs showed specific upregulated expression of totipotent genes enriched in immune response associated pathways **(Figure 3D)**. In summary, our analysis reveals differences in expression patterns of the totipotent signatures among 2CLCs, cTotiSCs, and TPSs, highlighting the potential of leveraging these *in vitro* models for exploring distinct transcriptomic features inherent to totipotency.

### Key histone modifications correlated with gene expression in 2CLCs

Our analysis showed that 2CLCs partially recapitulated the transcriptome features of totipotency, as evidenced by the extremely high expression of totipotent and minor ZGA genes in 2CLCs compared to other cells **(Figure 3B)**. To explore key epigenetic modifications modulating totipotent signatures in 2CLCs, we initially examined the correlation between gene expression and histone modifications, which were previously shown to regulate early development [54], including H3K4me3, H3K27ac, H3K27me3, and H3K79me3. When considering all protein-coding genes (PCGs) whose promoters regions overlap with histone modification peaks (± 5 kb around the transcription start site of each PCG), we only observed that H3K4me3 modification showed association with their expression **(Figure 4A)**. Also, compared to other types of histone modifications we analyzed, H3K4me3 illustrated much stronger association with both totipotent genes (Pearson’s correlation = 0.19, two-sided P-value = 1.12E-07) and minor ZGA genes (Pearson’s correlation = 0.23, two-sided P-value = 4.6E-03) **(Figure 4A, Table S4)**. In addition, the marker H3K27ac positively correlated with expression of both totipotent and minor ZGA genes **(Figures S1A and S1B)**. The Polycomb repression marker H3K27me3 [55] showed no correlation with expression of all PCGs, totipotent genes, and minor ZGA genes in 2CLCs **(Figures 4B and S1C)**, which is in line with the previous finding that H3K27me3 does not modulate 2C-like transition in mESCs [11]. H3K79me3 only showed a slightly positive association with totipotent genes but not all PCGs **(Figures 4B and S1D)**. Furthermore, comparing to H3K27ac, H3K27me3, and H3K79me3, we only observed that enriched H3K4me3 modification around the transcription start site (TSS) of the nearest PCGs in 2CLCs **(Figure 4C)**. Taken together, our example suggests that histone modifications, particularly H3K4me3, might play an important role in defining the molecular underpinning of totipotency in 2CLCs.

**Figure 4.**
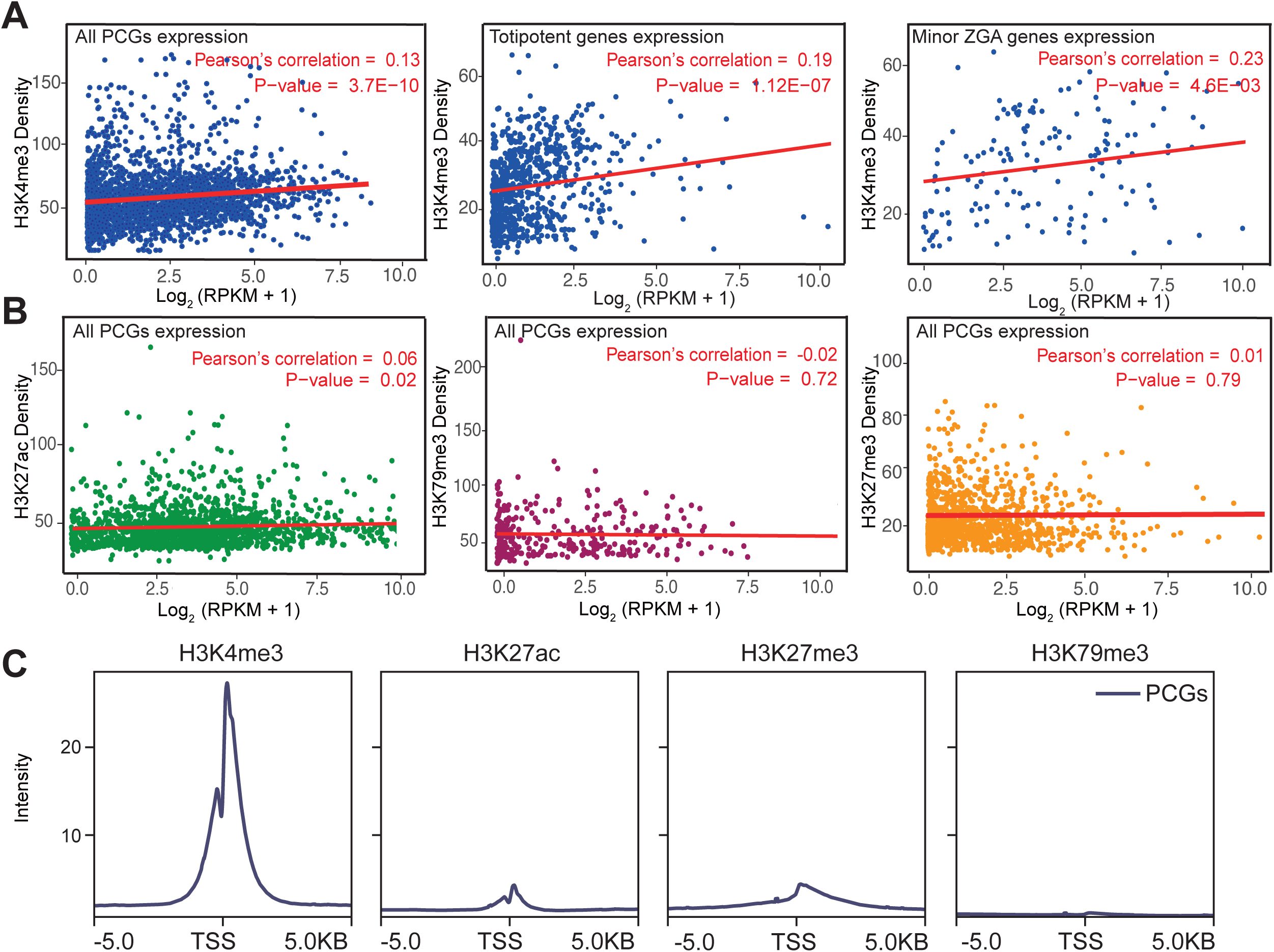
Identification of key histone modifications that modulate gene expression in 2CLCs. **A.** Correlation between H3K4me3 modification and expression of all protein-coding genes (PCGs; left), totipotent genes (middle), and minor ZGA genes (right). The x axis is normalized gene expression [log_2_ (RPKM +1)], and the y axis is the density of reads overlapping with H3K4me3 modified regions (± 5 kb around TSS) in 2CLCs. **B.** Correlation between expression of all protein-coding genes (PCGs) and H3K27ac (left), H3K79me3 (middle), H3K27me3 (right) modification. The x axis is normalized gene expression [log_2_ (RPKM +1)] and the y axis is the density of reads overlapping with histone modified regions (± 5 kb around TSS) and in 2CLCs. **C.** Density distribution of reads overlapping with histone modified regions (± 5 kb around TSS) in 2CLCs. The x axis is the distance from the transcription start site. And the y axis is the reads density.

### The potential role of H3K4me3-*Zscan4* axis in 2CLCs

*Zscan4*-family genes are typically totipotent signature genes in 2CLCs [46]. Next, we investigated gene expression and epigenetic regulatory patterns around *Zscan4a, Zscan4c, Zscan4d, and Zscan4f* genes in 2-cell embryos, selected *in vitro* TSCs models (2CLCs, TPSs, ciTotiSCs, and TLSCs), and pluripotent mESCs. *Zscan4c* only expressed in 2-cell embryos and *in vitro* TSCs models but not in mESCs **(Figure 5A)**. Open accessible regions around *Zscan4c* also show a similar totipotent-specific pattern with accessible peaks existing only in 2-cell embryos and 2CLCs, when compared to mESCs. Importantly, H3K4me3 peaks around *Zscan4c* were specifically presented in 2CLCs and 2-cell embryos but not in mESCs as well **(Figure 5A)**, suggesting the potential totipotency-relevant role of H3K4me3-*Zscan4c* axis. Similar expression and epigenetic modification patterns were also observed around *Zscan4a, Zscan4d*, and *Zscan4f* **(Figure S2).** These findings are consistent with previous studies [46, 56] showing totipotent-specific patterns both in expression and H3K4me3 modification surrounding *Zscan4* family genes.

**Figure 5.**
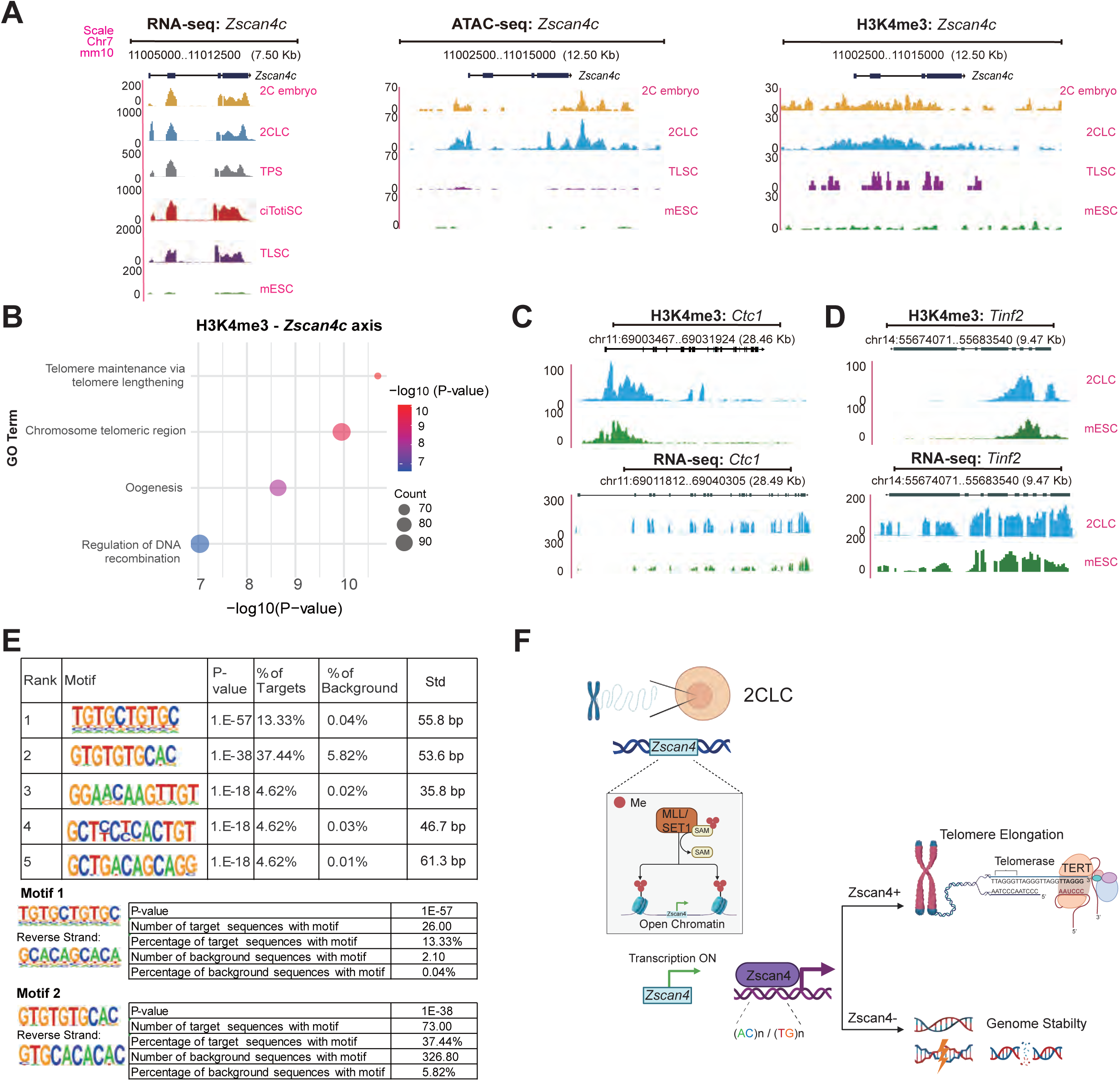
The potential role of H3K4me3-*Zscan4c* axis in 2CLCs. **A.** Reads coverage represents gene expression, chromatin accessibility, and H3K4me3 modifications (from left to right) around gene *Zscan4c* in 2C embryo, 2CLC, TPS, ciTotiSC, TLSC, and mESC. 2CLC: 2-cell-like cell. TPS: totipotent potential stem cell; ciTotiSC: chemically induced totipotent stem cell; TLSC: totipotent-like stem cell; mESC: mouse embryonic stem cell. **B.** Functional annotation (gene ontology) of H3K4me3-*Zscan4c* axis (H3K4me3 modified regions located within ± 2 kb around TSS of *Zscan4c*) in 2CLCs. The x axis is the -log_10_(two-sided P-value), and the y axis represents the enriched GO terms. **C.** Reads coverage represents H3K4me3 modification and gene expression around *Ctc1* in 2CLC and mESC. 2CLC: 2-cell-like cell. mESC: mouse embryonic stem cell. **D.** Reads coverage represents H3K4me3 modification and gene expression around *Tinf2* in 2CLC and mESC. 2CLC: 2-cell-like cell. mESC: mouse embryonic stem cell. **E.** Motif enrichment results for *Zscan4c* in *Zscan4c*-over-expressed mESCs. **F.** Schematic illustrating the putative role of *Zscan4c* in maintaining genomic stability and telomere elongation.

Next, we investigated the function role of H3K4me3-*Zscan4c* axis in 2CLCs using the ‘Pathway Enrichment’ module. Intriguingly, we found that H3K4me3-*Zscan4c* axis suggested an enrichment in telomere maintenance and chromosome telomeric region relevant pathways (**Figure 5B**). In addition to *Zscan4c*, we also observed that H3K4me3 modification and expression of other genes involved in telomere regulation and maintenance, such as *Ctc1* [57] and *Tinf2* [58], showed similar patterns of enrichment in 2CLCs compared to mESCs (**Figures 5C and 5D**). Altogether, these results indicate the potential role of H3K4me3-*Zscan4* axis in regulating the telomere in 2CLCs **(Figures 5B-5D; Table S5)**.

Furthermore, we examined the target sequences of *Zscan4c* in *Zscan4c*-over-expressed mESCs (ChIP-seq data in Toti) using the ‘Motif Enrichment’ module. The top 2 enriched binding motifs displayed a repeated CA/TG pattern, suggesting the microsatellite signatures **(Figure 5E)**. The (CA)n/(TG)n microsatellites are susceptible to chromosomal recombination, which affects the genome integrity [59, 60]. *Zscan4* was previously found to recognize and bind to (CA)n/(TG)n microsatellites in nucleosomes, preventing them from forming an energetically unfavorable structure known as Z-DNA and disassembling under torsional stress [61]. This mechanism potentially functions as a developmentally regulated safeguard that maintains genomic integrity in totipotent embryos, which is in line with the observed enrichment in ‘regulation of DNA recombination’ pathway for H3K4me3-*Zscan4c* axis illustrated in **Figure 5B**.

Collectively, our primary analysis suggests the hypothesized role of *Zscan4* in preserving genomic integrity by maintaining telomeres and safeguarding microsatellite regions in 2CLCs. As illustrated by this case study, we show that Toti could facilitate users to understand the potential function of H3K4me3 in regulating genomic stability in TSCs through activating *Zscan4* **(Figure 5F)**. Our case study exemplified how to apply Toti to explore discrepancies of transcriptome and epigenome nuances underlying totipotency across mESCs, 2-cell embryos, and *in vitro* TSC models.

## Discussion

To the best of our knowledge, Toti is the first and unique multi-omics database exclusively developed for investigation of transcriptional and epigenetic factors shaping totipotency and can be accessed through a user-friendly Web interface (http://toti.zju.edu.cn/). Toti unprecedentedly provides a comprehensive repository of transcriptomic (RNA-seq and scRNA-seq) and epigenomic (ATAC-seq, ChIP-seq, CUT&RUN, CUT&TAG and bisulfite sequencing) data from 4,671 human and 3,594 mouse samples, covering *in vivo*, *in vitro* and genome-edited embryonic TSCs, TSC-like cells, PSCs and embryos spanning preimplantation stages, setting it apart from other embryonic and stem-cell resources, such as dbEmbryo [44], DevOmics [43] and StemCellDB [31]. The ‘Search’ module enables users to easily and effectively focus on their interested studies/datasets by searching for genes, sequencing types or other relevant keywords. This ‘Browse’ module provides insights into the (dis)similarity across *in vivo* and *in vitro* totipotent cells, PSCs, and other embryonic cells in human and mouse by allowing intuitive and comparative visualization of epigenetic and transcriptomic features. With three sub-modules designed in the ‘Analysis’ section, Toti also facilitates users to prioritize key motifs and biological pathways that might be able to capture gene regulatory logic underlying totipotency. Furthermore, Toti allows an online scRNA-seq analysis module tailored to investigation on cellular transcriptome underpinning of the establishment and exit of totipotency at single-cell resolution. We conducted a case study to illustrate the potential application of Toti in exploiting totipotency research. While we acknowledge that the analyses in this case study may lack rigor, they sufficiently demonstrate the potential of Toti in this field. We believe Toti constitutes a comprehensive resource and opens new avenues for understanding the intricacies of totipotency.

In the future, we will continuously maintain and update Toti by curating and incorporating the latest findings in TSCs studies. As TSCs are the progenitors of all cell types in mammals and have great potential for regenerative medicine, we will continue to expand our collection of publicly available multi-omics data of TSCs to more species, such as pigs and monkeys. In addition, we will continue to integrate other data types, such as scATAC-seq and Hi-C data, to further improve our understanding of totipotency-associated regulatory networks. These additions are anticipated to provide insights into the molecular basis of totipotency and facilitate researchers to stabilize totipotency *in vitro*.

## Acknowledgements

This work is supported by the National Natural Science Foundation of China [ 32300534, 32270659 to X.S. and 32470840, 32270852 to X.F.], the Open Project of Jiangsu Provincial Science and Technology Resources (Clinical Resources) Coordination Service Platform [TC2023006 to X.S.], the Natural Science Foundation of Zhejiang Province [LZ24C120001 to X.F.], the Key R&D Program of Zhejiang Province [2024SSYS0021 to X.S. and 2024SSYS0020 to X.F.] and the hundred talents program of Zhejiang University to X.S. and X.F. We thank Dr. Jingfa Xiao, Dr. Yu Hou, Jinying Zhang, Tong Li, Pengcheng Wang, Dong Zou, and Yunna Li for their assistance in database construction. We thank the technical support from the Core Facilities, Liangzhu Laboratory, Zhejiang University.

## Author contributions

X.S., X.F., and Y.C. conceived, planned, and oversaw the study and wrote the manuscript. Y.C. and R.Z. performed the data analyses. X.S. and Y.C. developed the Web database. S.J., D.Z., S.C., X.F. and X.S. assisted with data collection and manuscript revision. All named authors read and approved the final manuscript.

