## Supplementary Figures for "Toti: an integrated multi-omics database to decipher the epigenetic regulation of gene expression in totipotent stem cells"

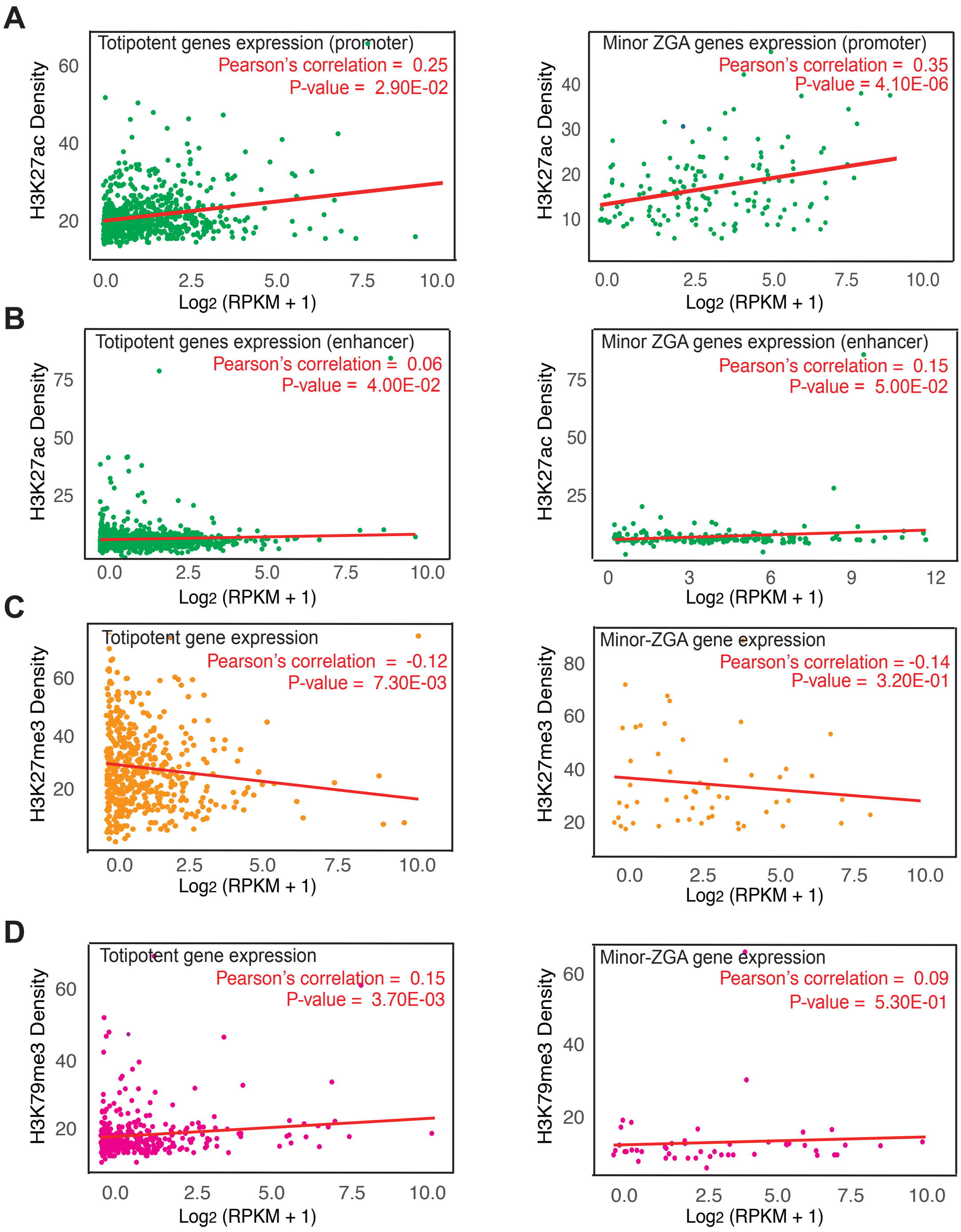


**Figure S1. Identification of key histone modifications that modulate gene expression in 2CLCs.** (A) Correlation between H3K27ac modification and expression of totipotent genes (left), minor ZGA genes (right). The x axis is normalized gene expression [$\log_2$ (RPKM +1)], and the y axis is the density of reads overlapping with H3K27ac modified regions (± 5 kb around TSS) in 2CLCs. (B) Correlation between H3K27ac modification and expression of totipotent genes (left) and minor ZGA genes (right). The x axis is normalized gene expression [$\log_2$ (RPKM +1)], and the y axis is the density of reads overlapping with H3K27ac modified regions (± 4 Mb around TSS) in 2CLCs. (C) Correlation between H3K27me3 modification and expression of totipotent genes (left) and minor ZGA genes (right). The x axis is normalized gene expression [$\log_2$ (RPKM +1)], and the y axis is the density of reads overlapping with H3K27me3 modified regions (± 5 kb around TSS) in 2CLCs. (D) Correlation between H3K27me3 modification and expression of totipotent genes (left) and minor ZGA genes (right). The x axis is normalized gene expression [$\log_2$ (RPKM +1)], and the y axis is the density of reads overlapping with H3K79me3 modified regions (± 5 kb around TSS) in 2CLCs.

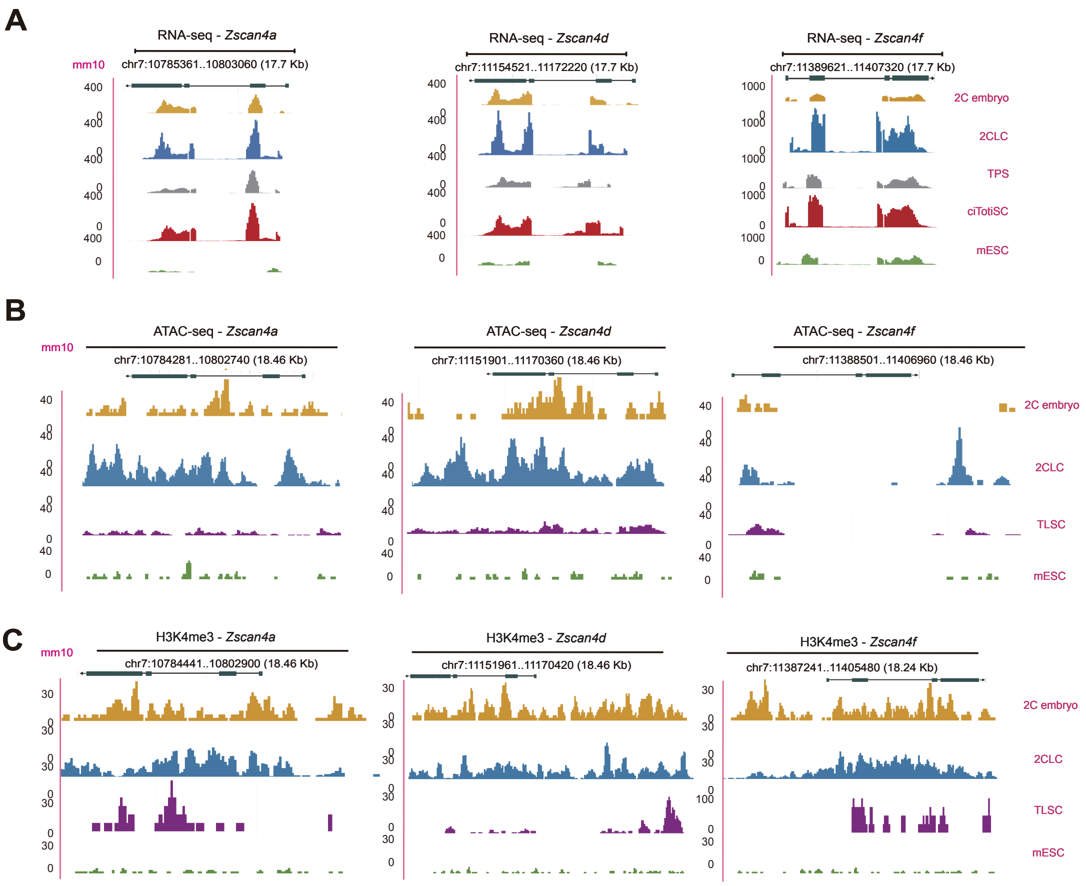


**Figure S2. Reads distribution.** (A) Reads coverage represents gene expression, (B) chromatin accessibility, and (C) H3K4me3 modification around gene *Zscan4a* (left), *Zscan4d* (middle), and *Zscan4f* (right), in 2C embryo, 2CLC, TLSC, and mESC. 2CLC: 2-cell-like cell, TLSC: totipotent-like stem cell, mESC: mouse embryonic stem cell.

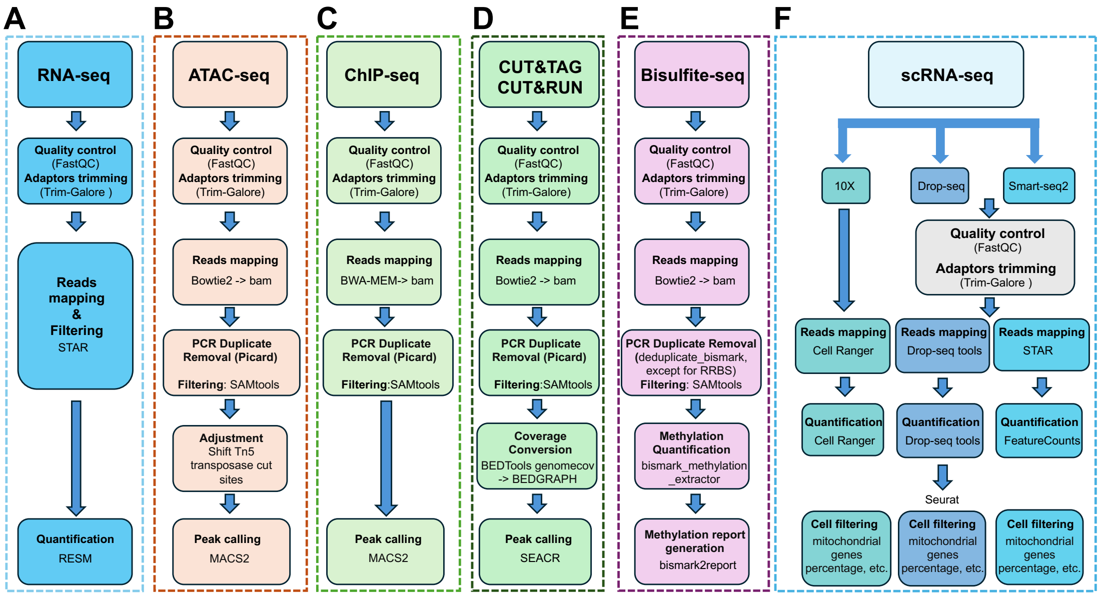


**Figure S3. Data analysis pipeline.** (A) RNA-seq, (B) ATAC-seq, (C) ChIP-seq, (D) CUT&TAG, CUT&RUN, (E) Bisulfite-seq, and (F) scRNA-seq.
